# Algorithmic Parameter Estimation and Uncertainty Quantification for Hodgkin-Huxley Neuron Models

**DOI:** 10.1101/2021.11.18.469189

**Authors:** Y. Curtis Wang, Nirvik Sinha, Johann Rudi, James Velasco, Gideon Idumah, Randall K. Powers, Charles J. Heckman, Matthieu Chardon

## Abstract

Experimental data-based parameter search for Hodgkin–Huxley-style (HH) neuron models is a major challenge for neuroscientists and neuroengineers. Current search strategies are often computationally expensive, are slow to converge, have difficulty handling nonlinearities or multimodalities in the objective function, or require good initial parameter guesses. Most important, many existing approaches lack quantification of uncertainties in parameter estimates even though such uncertainties are of immense biological significance. We propose a novel method for parameter inference and uncertainty quantification in a Bayesian framework using the Markov chain Monte Carlo (MCMC) approach. This approach incorporates prior knowledge about model parameters (as probability distributions) and aims to map the prior to a posterior distribution of parameters informed by both the model and the data. Furthermore, using the adaptive parallel tempering strategy for MCMC, we tackle the highly nonlinear, noisy, and multimodal loss function, which depends on the HH neuron model. We tested the robustness of our approach using the voltage trace data generated from a 9-parameter HH model using five levels of injected currents (0.0, 0.1, 0.2, 0.3, and 0.4 nA). Each test consisted of running the ground truth with its respective currents to estimate the model parameters. To simulate the condition for fitting a frequency-current (F-I) curve, we also introduced an aggregate objective that runs MCMC against all five levels simultaneously. We found that MCMC was able to produce many solutions with acceptable loss values (e.g., for 0.0 nA, 889 solutions were within 0.5% of the best solution and 1,595 solutions within 1% of the best solution). Thus, an adaptive parallel tempering MCMC search provides a “landscape” of the possible parameter sets with acceptable loss values in a tractable manner. Our approach is able to obtain an intelligently sampled global view of the solution distributions within a search range in a single computation. Additionally, the advantage of uncertainty quantification allows for exploration of further solution spaces, which can serve to better inform future experiments.

## Introduction

### Systematic parameter exploration: an unmet need for model building in neuro-science

#### Can we measure everything needed to build a model?

Complete specification of models in neuroscience or systems physiology requires identification of several parameters. For single neurons, these typically include geometric and electrical properties of the cell body, dendrites, and axons. At the network level, one needs to additionally specify the synaptic connectivity patterns along with the temporal and spatial properties of individual synapses. Even with the sophistication of modern experimental techniques, however, measuring all the necessary parameters is almost always impossible. Additionally, many single-neuron and network-level properties exhibit remarkable context-dependent variability, even within the same animal. For example, changes in the neuromodulatory environment alter the spiking dynamics of spinal motoneurons (***Heckman et al., 2009***) and thalamocortical neurons (***Pape and McCormick, 1989***) by influencing their ionic conductances. Likewise, changes mediated by long-term potentiation or depression at individual synapses propagate selectively throughout the network (***Fitzsimonds et al., 1997***). Thus, any single measurement at one time instance would be insufficient to inform all single-neuron/network properties because it cannot account for the wide spectrum of behaviors observed in vivo.

To complicate matters further, redundancy in biological systems leads to similar activity profiles that can be produced by many different neurons or neuronal networks with dissimilar properties (***Schulz et al., 2007***; ***Roffman et al., 2012***; ***Swensen and Bean, 2005***). For example, lateral pyloric neurons in the crab stomatogastric ganglion exhibit as much as two- to fourfold interanimal variability in three different ion channel densities and their corresponding mRNA levels (***Schulz et al., 2006***). The stomatogastric ganglia can also exhibit identical network activity from widely disparate synaptic strengths and neuron properties (***Prinz et al., 2004***). Therefore, even averaging a parameter from multiple experimental preparations may fail to generate the desired behavior in the computational models constructed from them (***Golowasch et al., 2002***). Systematic exploration of the parameter space is obligatory to fit a neuron or a neuronal network model to experimental data. Additionally, this systematic exploration may reveal deeper insights and motivate future experiments by unraveling undiscovered parameter combinations that might reproduce the same experimentally observed behavior.

#### Is manual tuning of model parameters still practicable?

Fitting experimental data to neuron models has been a major challenge for neuroscientists. In fact, the parameters of the first biologically realistic quantitative description of the neuronal membrane were hand fitted by ***Hodgkin and Huxley*** (***1952***). This approach has remained popular; for example, the peak conductances of ion channels in a model of an elemental leech heartbeat oscillator were hand tuned by ***Nadim et al.*** (***1995***). Likewise, both the single compartment and network parameters of a small group of neurons used to model the crustacean pyloric network were hand tuned by ***Soto-Treviño et al.*** (***2005***) to reproduce a variety of experimentally observed behaviors. However, the ever-increasing dimensionality and complexity of the neuron models (Hodgkin and Huxley’s description contained only three membrane currents), accompanied by the concomitant increase in computational resources available at the disposal of neuroscientists, have made hand tuning practically obsolete. Besides, hand tuning of parameters also introduces experimenter bias, because different parameters are assigned preconceived roles during the tuning process (***Van Geit et al., 2008***). Nevertheless, hand tuning to a certain extent may be unavoidable because the initial range of values of different parameters over which the automated search is to be performed must be determined by the experimenter based on physiological constraints.

### Computational inference for neural models

The body of research on parameter estimation for models of neural dynamics for single cells or circuits spans across various scientific communities, approaches, and neuron models. A gap exists, however, between researchers with rich physiological knowledge, on the one hand, and researchers working on new solution algorithms for inference, on the other hand. This gap presents opportunities to create a bridge between physiologists, engineers, computational scientists, and mathematicians by exploring existing and developing new inference techniques for neuron dynamics of single cells and circuits.

#### Multiple realizability: the dilemma of many truths

Single-neuron and network-level activities that are amenable to easy measurement (e.g., spike trains, voltage traces, local field potentials) can often be identical for different parameter combinations. Just as this situation renders experimental determination of parameter values infeasible, it also poses an incredible challenge to the computational neuroscientist trying to systematically infer a model from the experimental data. For example, ***Prinz et al.*** (***2004***) found indistinguishable network activity from widely disparate deterministic models. Similarly, ***Amarasingham et al.*** (***2015***) reported statistically indistinguishable spike train outputs for different statistical processes that model the firing rate of networks. ***Hartoyo et al.*** (***2019***) stressed that, for models of dynamical systems, very different parameter combinations can generate similar predictions/outputs. Additionally, the authors showed that sensitivities of predicted parameters can exhibit wide variability, leading to the conclusion that modeling in neuroscience is confronted by the major challenge of identifiability of model parameters. The demonstrative model used in our present work shows a bursting neuron producing similar neural activities from multiple sets of conductance densities (***Alonso and Marder, 2019***).

#### Previous approaches

Computational approaches for estimating parameters of neuron models based on ordinary differential equations (ODEs) include brute-force grid search (***Prinz et al., 2003***) as well as more advanced techniques using heuristics or trial-and-error approaches, which may consist of intricate sequences of regression steps (***Achard and De Schutter, 2006***; ***Van Geit et al., 2007***, ***2008***; ***Buhry et al., 2011***). The latter include simulated annealing, differential evolution, and genetic algorithms. These approaches have the disadvantage of slowly converging to an optimal set of parameters and thus being computationally expensive. Using gradient-based optimization to accelerate convergence (***Doi et al., 2002***; ***Toth et al., 2011***; ***Meliza et al., 2014***), on the other hand, can suffer from the nonconvexity and the strong nonlinearities in the objective manifold and require good initial guesses in order not to get stuck in local minima (thereby resulting in suboptimal parameter sets). Another shortcoming of recovering only one set of parameters (also called point estimates) is the lack of knowledge about the uncertainties or error bounds for the inferred parameters.

Attempts at using machine learning techniques based on artificial neural networks (ANNs) have recently shown promising results. For instance, in the context of the FitzHugh–Nagumo model (***Rudi et al., 2021***), an ANN was constructed and optimized to generate an inverse map that is able to predict model parameters from observational data. ***Bittner et al.*** (***2019***) developed generative models from deep learning. Retrieving associated uncertainties for inferred parameters has been performed by ***Gonçalves et al.*** (***2020***), using normalizing flows with Gaussian mixtures in order to generate approximations of posterior distributions for Hodgkin–Huxley-based inverse problems. However, machine learning techniques still suffer from the cost of generating large sets of data for training the artificial neural networks, where each training sample requires the numerical solution of a differential equation.

Theoretical neuroscientists have been developing and utilizing statistical methods for inference and uncertainty quantification (***Van Geit et al., 2007***; ***Vavoulis et al., 2012***) and have relied on Bayesian inference frameworks (***Ahmadian et al., 2011***; ***Doruk and Abosharb, 2019***) in order to estimate parameters and quantify uncertainties of the recovered parameters. The uncertainties in recovered parameters with Bayesian inference frameworks are represented by posterior density functions, which describe a “landscape” of more likely parameters (“peaks in the landscape”) and less likely parameters (“valleys in the landscape”; see Methods for details). Bayesian likelihoods or their approximations are constructed, which in turn enables direct maps to posterior densities without numerical solutions of neuron models (i.e., differential equations). ***Chen*** (***2013***) provides an overview of Bayesian methods for neural spike train analysis. Recently, ***René et al.*** (***2020***) and ***Schmutz et al.*** (***2020***) have utilized Markov chain Monte Carlo for inference from spike train data of population models.

### Using Markov chain Monte Carlo algorithms for inference

MCMC algorithms play a critical role in our approach for parameter estimation. For a brief introduction and background information about MCMC algorithms, we refer the reader to the Methods section.

#### Benefits and limitations of using MCMC in parameter searches

One of the main benefits of using MCMC is the detailed picture of the posterior “landscape” that it can provide. By analyzing the posterior, the uncertainty in the inferred parameters can be quantified, and the dependencies between different parameters in the model can be identified. Another benefit is that MCMC does not require derivatives of neuron models or the loss function for misfits of the data and model output. This is important in the case of the complex model and nondifferentiable loss function proposed by ***Alonso and Marder*** (***2019***), which we also use in the present work. Furthermore, the key advantage of parallel tempering MCMC is its ability to expose multimodality in the posterior (see Methods section for more details).

Arguably, possible limitations of MCMC may arise if the search space for the parameters becomes too large; that is, if too many parameters in the model need to be inferred. With increasing dimensions of the parameter space, increasingly more iterations of MCMC may be required, eventually leading to the algorithm becoming computationally prohibitive. In the inference that we are targeting in this work, the number of inferred parameters remains at amounts where these limitations do not occur.

#### Applications of MCMC in biological modeling

MCMC-based methods have generally been a popular choice for solving inverse problems in a statistical framework, where one is interested in uncertainties in addition to optimal solutions (***Smith, 2013***). In the context of dynamic systems in biology, MCMC techniques have been successfully employed, as described in review articles by ***Ballnus et al.*** (***2017***) and ***Valderrama-Bahamo*** (***2019***). Moreover, tempering MCMC methods have been shown to recover the multimodality of solutions in systems biology (***Caranica et al., 2018***; ***Gupta et al., 2020***). However, in the context of neural dynamics with complex Hodgkin–Huxley models, such as in ***Alonso and Marder*** (***2019***), which we target in this work, MCMC methods have not been attempted. The reason may be the computational challenges described above, for instance, multimodal posteriors.

### Objectives and contributions of this work

In this study we address the issue of multiple realizability of models in neuroscience. Specifically, we target the computational inference for HH-based neuron models with a state-of-the-art MCMC algorithm. Our goal is to estimate the parameters in such models and quantify their uncertainties.

We design the parameter estimation problem in a Bayesian framework: (i) parameter sets in a model are treated as probability distributions rather than one single optimal set; and (ii) the distributions of parameter sets can exhibit dependencies between individual values of the set, rather than assuming each parameter to be independent from the others. This Bayesian framework for parameter estimation does not assume that a single independent optimum can be reached. The likelihood is given implicitly in the form of a loss function between observational data and model output, hence requiring the numerical solution of the model for each new set of parameters. As a solution algorithm we utilize an extension MCMC, the parallel tempering MCMC method, which is critical for recovering distributions of parameter sets that are multimodal (i.e., multiple sets of parameters of a neuron model are recovered for a single set of observational data). Further, we visualize solution maps of the multimodal posterior that enable us to quantify which parameters can reproduce the observational data more closely.

## Results

The results of our study show that our design of the inverse problem combined with parallel tempering MCMC for solving the inverse problem is able to successfully overcome the inference challenges by using a systematic approach. This MCMC algorithm can recover multiple parameter sets where data and model output are consistent. The algorithm requires only an initial setup of the prior information (i.e., assumptions on the distributions of the parameter values), and it especially does not require manual intervention, such as for widening or narrowing bounds for parameter values, during the search. Parallel tempering MCMC returns distributions of sets of parameters, which we refer to as *solution maps* of the posterior. With these solution maps physiologists can investigate the results from MCMC in order to quickly decide which parameter sets are indeed relevant from a physiological perspective. This is a main advantage of *solution maps* compared with *solution points* (i.e., a single set of “optimal” parameters) obtained from optimization algorithms.

### Inference proof of concept with a Hodgkin–Huxley model

We illustrate the results of the parallel tempering MCMC method in a simple inverse problem setup, which nevertheless exhibits the difficulties of more complex inference setups in neural dynamics. We use a relatively simple and well-known HH model (***Hodgkin and Huxley, 1952***; ***Mainen et al., 1995***) and the commonly used mean squared error (MSE) as the loss function to measure the fit between data and model outputs. To set up the inverse problem, we simulate the voltage trace for a given set of sodium and potassium conductances, denoted as the “true” parameter set, which is *g_Na_* = 200 *pS*/*μm*^2^ for sodium and *g_K_* = 50 *pS*/*μm*^2^ for potassium. This voltage trace serves as the data for our inverse problem.

The first goal of the inverse problem is to recover the true sodium and potassium conductances. To demonstrate the challenges of the problem, we visualize the loss function that would have to be minimized to find the optimum in ***Figure 1***A, where the colors indicate the (positive) loss value. The true parameter set is located in a “valley” in the loss “landscape,” whereas the loss is large for parameters that produce large discrepancies between model output and data such as shown in graph S1 corresponding to point S1 in ***Figure 1***A. However, multiple sets of parameters exist that are local minima in this landscape, and these multiple local minima pose major challenges because optimization algorithms may present one of them as the “optimal” solution. These local minima will not give a sufficiently good fit of model outputs vs. data, as is illustrated in graph S2 of ***Figure 1*** that corresponds to point S2 in the landscape (***Figure 1***A). Such local minima are known to be problematic for numerical optimization algorithms (***Nocedal and Wright, 2006***).

**Figure 1.**
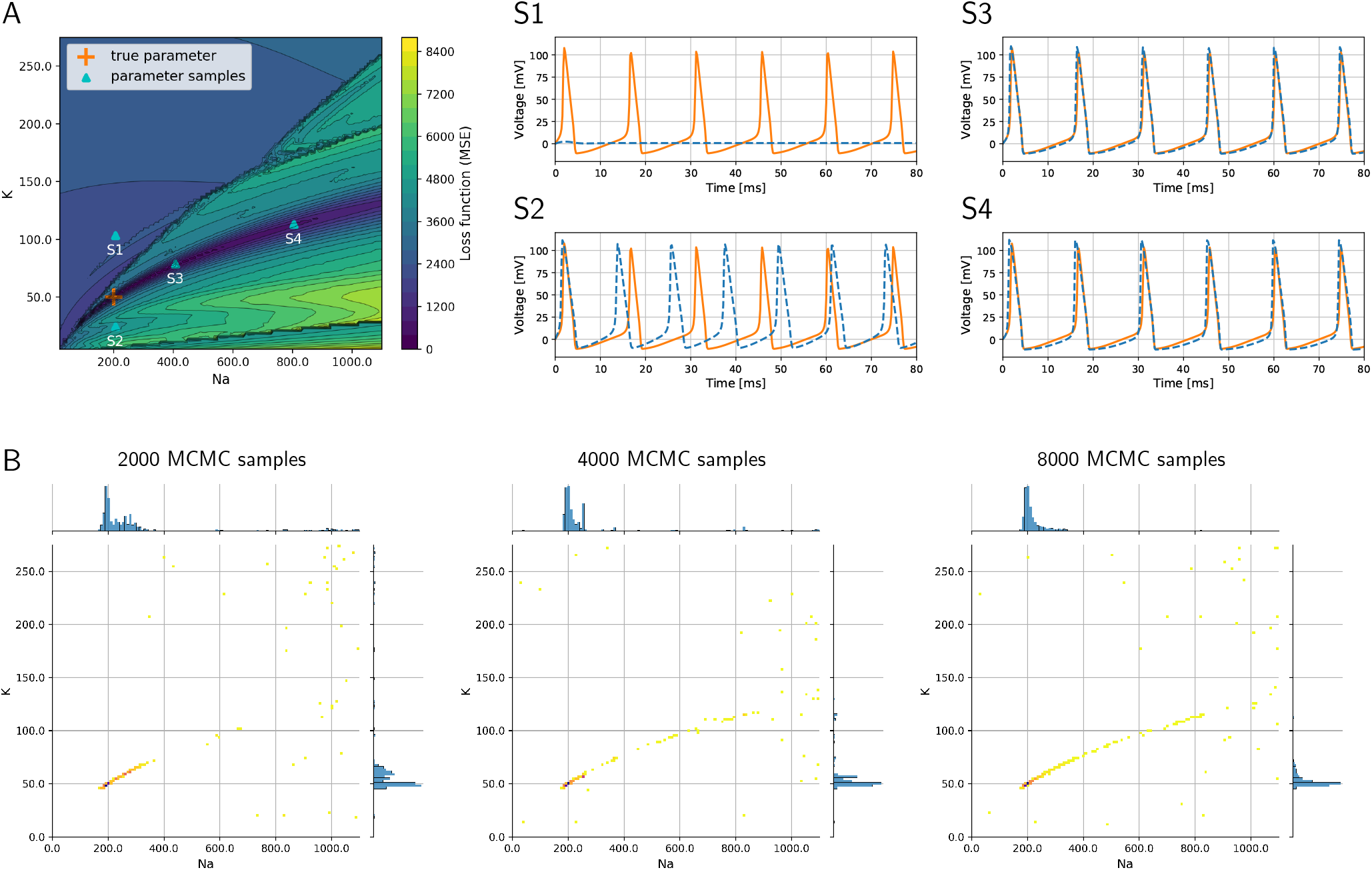
Loss function corresponding to the inverse problem with a simple HH model and how MCMC is able to successfully recover optimal parameters and quantify uncertainties and parameter trade-offs. A: Loss function landscape (colors) over the parameters, sodium (*g_Na_*) and (*g_K_*) potassium conductances. The “true” parameter set represents a global minimizer of the loss; the valley of the loss along points S3 and S4 (in dark blue color of loss) translates to parameter uncertainties (or trade-offs). Graphs S1–S4 depict voltage traces at corresponding points in A. B: Densities of MCMC samples, where the dark purple color of highest density overlaps with the true parameter set. As the number of MCMC samples increases, the method recovers the valley of the loss landscape A and hence quantifies uncertainties in the parameters.

The second goal of the inverse problem is to, additionally, quantify the uncertainty with respect to these parameters and understand the sensitivities of the parameters. For instance, in ***Figure 1***, the graphs S3 and S4 illustrate how different parameters produce voltage traces similar to the data, while at the same time this uncertainty in the parameters is visible in the landscape (***Figure 1***A) as a valley containing points S3 and S4. In order to recover these uncertainties, the formulation of the inverse problem in a Bayesian framework becomes critical.

The numerical solution of the Bayesian inverse problem with parallel tempering MCMC successfully provides a probability density spanned by the two-dimensional parameter space. The density is large, where the loss function in ***Figure 1***A has its major valley. We show the progress of the MCMC algorithm in ***Figure 1***B as the number of collected samples increases from 2,000 to 8,000. As the sample count (i.e., iteration count of MCMC) grows, the algorithm generates a longer tail along the valley, showing a clearer picture of the parameter uncertainties. The true parameter values are clearly visible in the high-density region as a dark blue area around 200 and 50 *pS*/*μm*^2^ for sodium and potassium conductances, respectively. Furthermore, the tail to the upper right of the true parameter set shows the trade-off between sodium and potassium, when model outputs keep being consistent with data even though the values of the parameters deviate from the truth.

These results serve as a proof of concept that the parallel tempering MCMC algorithm can successfully tackle multimodal losses. Next we transition to a more complex model for neural dynamics, which will be the main result of this paper.

### Inference with the complex Alonso–Marder (AM) model

#### AM model – Posterior distribution

MCMC samples the posterior distribution to generate a solution map of the posterior. Since we have nine uncertain parameters that we want to infer, the posterior is a distribution in a nine-dimensional space. To visualize the nine-dimensional space, we consider one or two parameters at a time, where the remaining parameter dimensions are marginalized (i.e., summed up). The plots in ***Figure 2*** show the solution map of the posterior. The denser regions within ***Figure 2*** are parameter value sets that return lower loss function values; therefore they represent better fits between data and model outputs.

**Figure 2.**
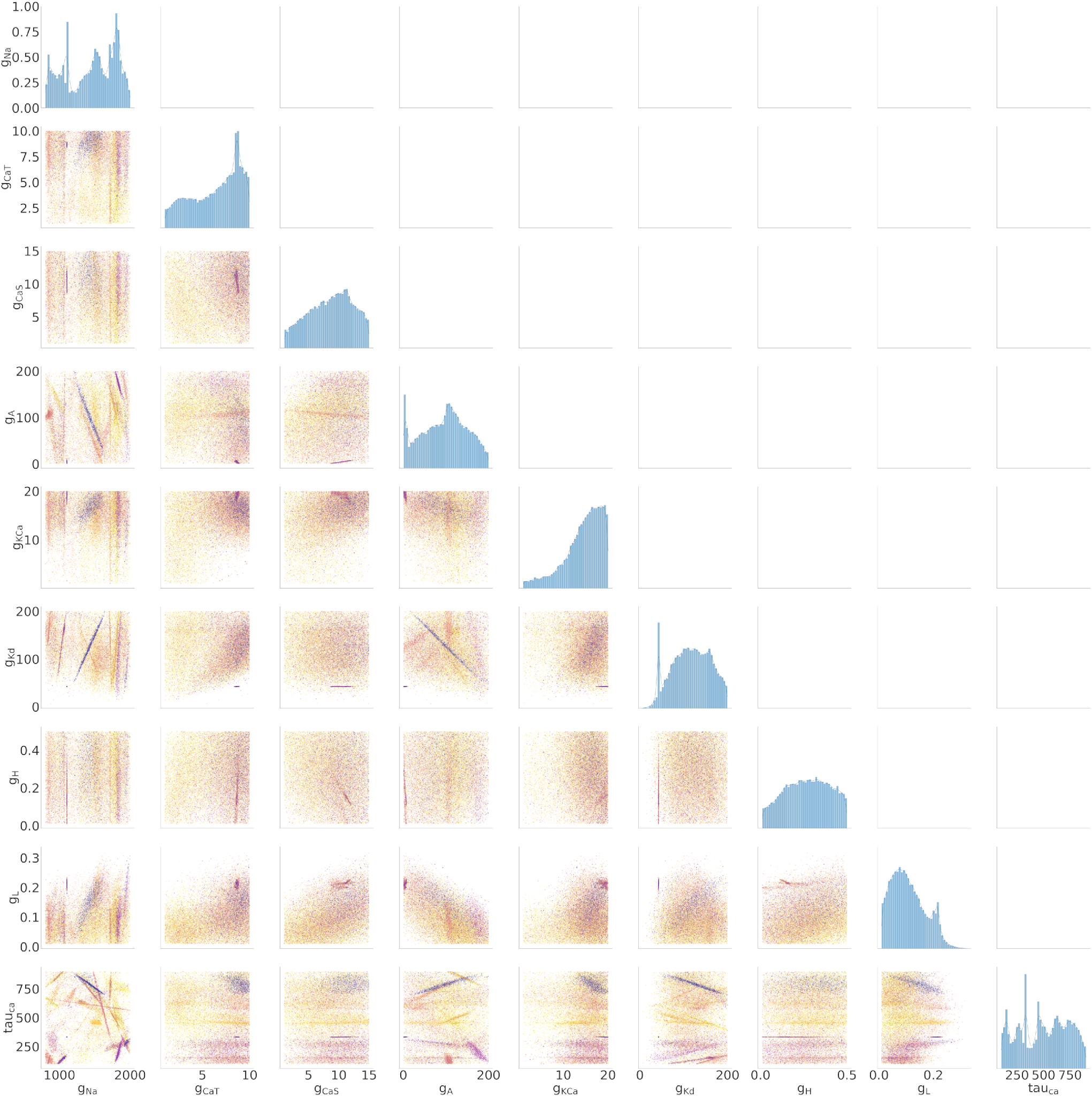
Final solution map of the posterior for the 9 parameters of the AM model for the **individual constraint** of the 0.0 nA injected current. The diagonal plots show the 1D marginals of each parameter. The x-axis is the search range of the parameter, and the y-axis is the normalized distribution. The remaining plots are the 2D marginals of pairs of parameters. The x- and y-axes are the range of the pairs of parameters. Each point within each plot is a sample found by the MCMC algorithm. The colors indicate the associated loss value of this sample, where the purple color indicates a lower loss value and the yellow color indicates a higher loss value. Density of points will contribute to the darkness of an area. The more pronounced a color signifies that more points are overlaid on top of each other. For example, in the top left 2D marginal [*g_CaT_, g_Na_*] the dark purple line around [*g_CaT_, g_Na_*] ~ [9, 1200] is an accumulation of many overlaid points. This density can also be traced back to the 1D marginals to the top and the left side.

Along the diagonal of ***Figure 2*** are the histograms for the (1D marginal) distribution for each parameter. For the 2D marginals in the lower triangle of ***Figure 2***, the points have the opacity set to darker where their density is higher. This setup allows for easier visualization of trends within the posterior distribution with regard to parameter values. Note that the denser points for a particular parameter (*g_Na_*, for example) are not necessarily in the denser regions for another parameter (*g_Kd_*, for example), if one considers different plots. Such links between two parameters can be established only when *g_Na_*, *g_Kd_* are plotted along the two axes of the same 2D marginal.

#### AM model – Individual constraint

We ran MCMC sampling on the AM model using individualized currents (0.0 to 0.4 nA at increments of 0.1 nA). This dataset is called the “Individual Constraint.” We also ran the MCMC search as a single aggregated search with all currents at once, which is discussed in a subsequent section. Below we present results from the individual constraints.

The MCMC search produced solution maps of the posteriors for each of the individual currents. Within each of the maps is a parameter set with the lowest loss value ***Table 1***. In the traditional thinking of optimization, this set can be considered to be the optimal solution.

**Table 1.** Parameter sets for the lowest loss values calculated by the MCMC algorithm. The range of the parameter priors is also provided in the first row. The units for the conductances are in *pS*/*μm*^2^.

| Injected current (nA) | $g_{Na}$ | $g_{CaT}$ | $g_{CaS}$ | $g_A$ | $g_{KCa}$ | $g_{Kd}$ | $g_H$ | $g_L$ | $\tau_{ca}$ | Loss |
| --- | --- | --- | --- | --- | --- | --- | --- | --- | --- | --- |
| uniform priors | [800, 2000] | [1, 10] | [1, 15] | [1, 200] | [1, 20] | [1, 200] | [0.01, 0.5] | [0.01, 0.5] | [100, 900] | – |
| 0.0 | 1484.10 | 9.54 | 11.07 | 78.73 | 18.13 | 144.03 | 0.027 | 0.119 | 767.54 | 0.0008 |
| 0.1 | 1543.74 | 9.90 | 14.27 | 73.17 | 19.74 | 171.16 | 0.067 | 0.149 | 768.77 | 0.0002 |
| 0.2 | 1768.69 | 6.64 | 13.52 | 118.36 | 18.01 | 68.42 | 0.096 | 0.075 | 739.20 | 0.0348 |
| 0.3 | 1461.32 | 6.63 | 10.14 | 86.07 | 10.41 | 62.39 | 0.425 | 0.193 | 452.63 | 0.0013 |
| 0.4 | 1973.44 | 4.06 | 9.93 | 168.56 | 1.24 | 29.14 | 0.221 | 0.183 | 206.94 | 0.0047 |
| aggregate | 1335.84 | 9.18 | 14.97 | 102.72 | 18.84 | 142.91 | 0.363 | 0.163 | 802.44 | 8.8109 |
| ground truth | 1076.39 | 6.41 | 10.05 | 8.04 | 17.58 | 124.09 | 0.113 | 0.176 | 653.50 | – |

In addition to producing a series of parameter sets with low loss values, the MCMC algorithm is able to produce the loss function map (posterior distribution) for all 9 parameters of the model within the search range and injected currents (0.0 to 0.4 nA, 0.1 nA increments).

We summarized these findings in a large arrangement of plots in ***Figure 2*** for 0.0 nA infected current and for currents 0.1–0.4 in the Appendix. The rows and columns of subplots are each associated with one of the parameters (*g_Na_*, *g_CaT_*, *g_CaS_*, *g_A_*, *g_KCa_*, *g_Kd_*, *g_H_*, *g_L_*, and *τ_Ca_*). As such, ***Figure 2*** resembles a triangular matrix, where the upper right triangle is redundant because it is symmetric to the lower left triangle.

##### 1D Marginals

The diagonal portion of the subplot matrix shows the one-dimensional marginals (distribution histograms) of the solutions for each of the individual parameters. The x-axis represents the range of values, and the y-axis represents the probability of that value producing a good fit between data and model outputs. These distributions provide both the likeliest solutions (i.e., the peaks) as well as their uncertainty (distribution around the peaks). For example, the top left 1D marginal of the sodium conductance *g_Na_* has 4 peaks. From those peaks, one can quickly quantify the uncertainty of each of those probable solutions.

##### 2D Marginals

The remaining subplots are the two-dimensional marginals of the posterior distributions. Each row and column represent a parameter *θ* of the model, and each of the points within the subfigure is a solution found by the MCMC algorithm. To show the probability distribution within the solution space, we set the alpha value (the transparency of the color) of each solution to *α* = 10^−3^ (where opaque is *α* = 1). A highly probable region has many more points and therefore will appear more opaque. The color of each point represents the loss value ranging from blue to yellow (low to high).

These plots show the dependence of each parameter on each other. Some of these dependencies are linear (for example, [row, col]: [*g_Kd_*, *g_Na_*]), and some are nonlinear (for example, [row, col]: [*g_Kd_*, *g_CaT_*]). Some of the linear dependencies are vertical and horizontal, signifying that one parameter is independent from the other (for example, vertical [*g_Kd_*, *g_CaT_*], [*g_H_*, *g_Na_*] and horizontal [*τ_Ca_*, *g_Cas_*]).

#### AM model – Aggregate constraint

To test the versatility of the MCMC algorithm at solving the inverse problem for HH-type equations, we combined all the input currents into a single analysis called “aggregate constraint” analysis (more details given in Methods).

Just as shown for the individual constraints, the aggregate constraint also have a solution map of the posterior. This map is shown in ***Figure 3*** where the 1D marginals and 2D marginals are illustrated. The same analysis can be performed to determine the peaks and variations of individual parameters using the 1D marginals and the dependence of two parameters using the 2D marginals. Compared with the individual constraint, the loss values are higher; hence the yellow colors dominate in these maps.

**Figure 3.**
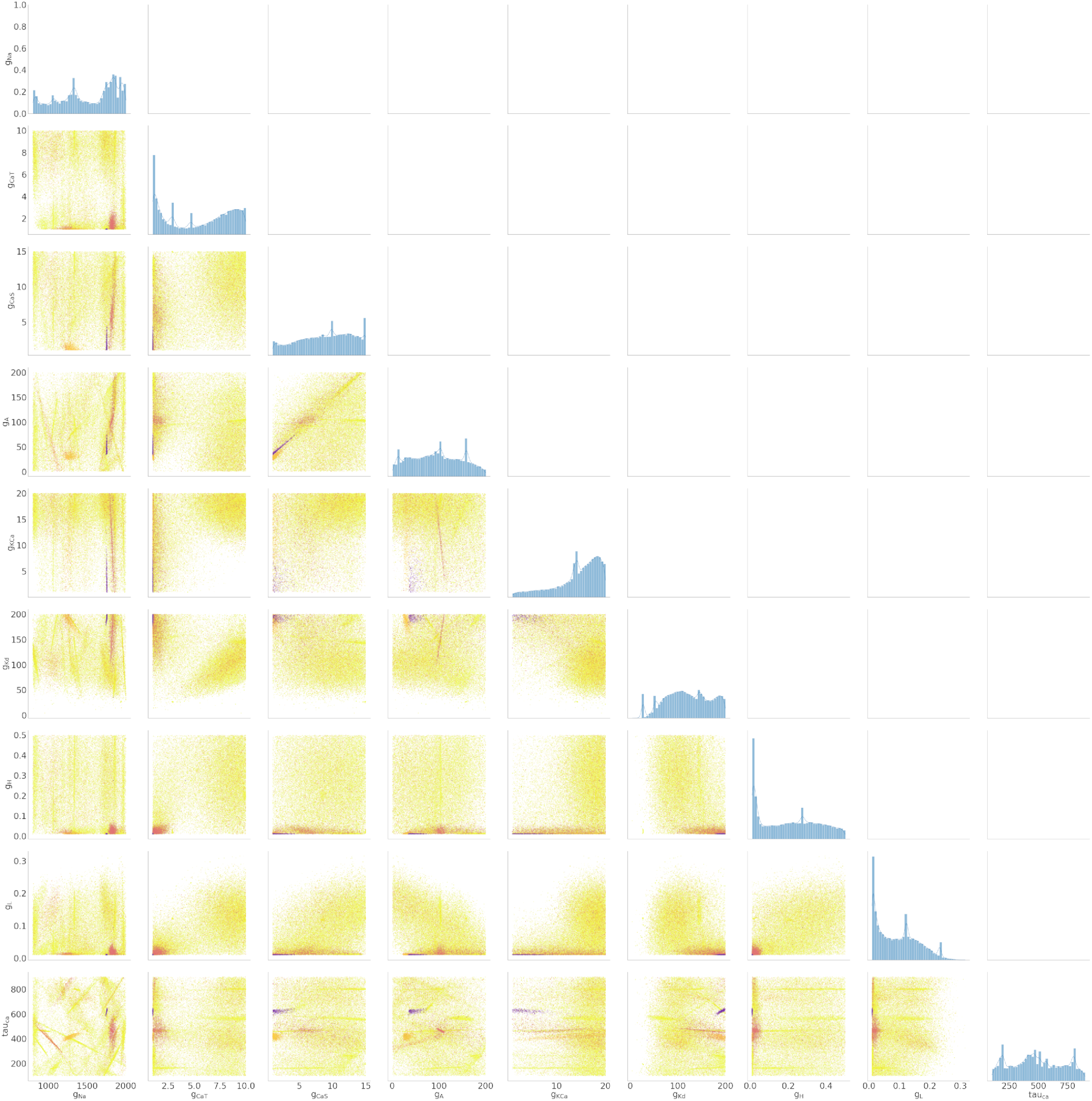
Solution maps of the posterior for the 9 parameters of the AM model for the **aggregate constraint**. The diagonal plots show the 1D marginals of each parameter. The x-axis is the search range of the parameter, and the y-axis is the normalized distribution. The remaining plots are the 2D marginals of pairs of parameters. The x- and y-axes are the range of the pairs of parameters. Each point within each plot is a sample found by the MCMC algorithm. The colors indicate the associated loss value of this sample, where purple color indicates a lower loss value and yellow color indicates a higher loss value. The density of the points will contribute to the darkness of an area. More pronounced color signifies that more points are overlaid on top of each other. This density can also be traced back to the 1D marginals to the top and the left side.

We found that the aggregate constraint solution map encompasses a subset of the individual constraint solution maps. To illustrate this, we show the intersection (multiplication of the posterior) of the kernel density estimates (KDEs) of the solution maps from the aggregate constraint and the two individual constraints 0.0 nA and 0.2 nA as an example in ***Figure 4***. First we calculated the intersection between the two individual constraints to extract the common set of solutions between the two. This first intersection can be interpreted as the solution map if only 0.0 nA and 0.2 nA were aggregated. We then calculated the intersection of the aggregate map with the latter solution.

**Figure 4.**
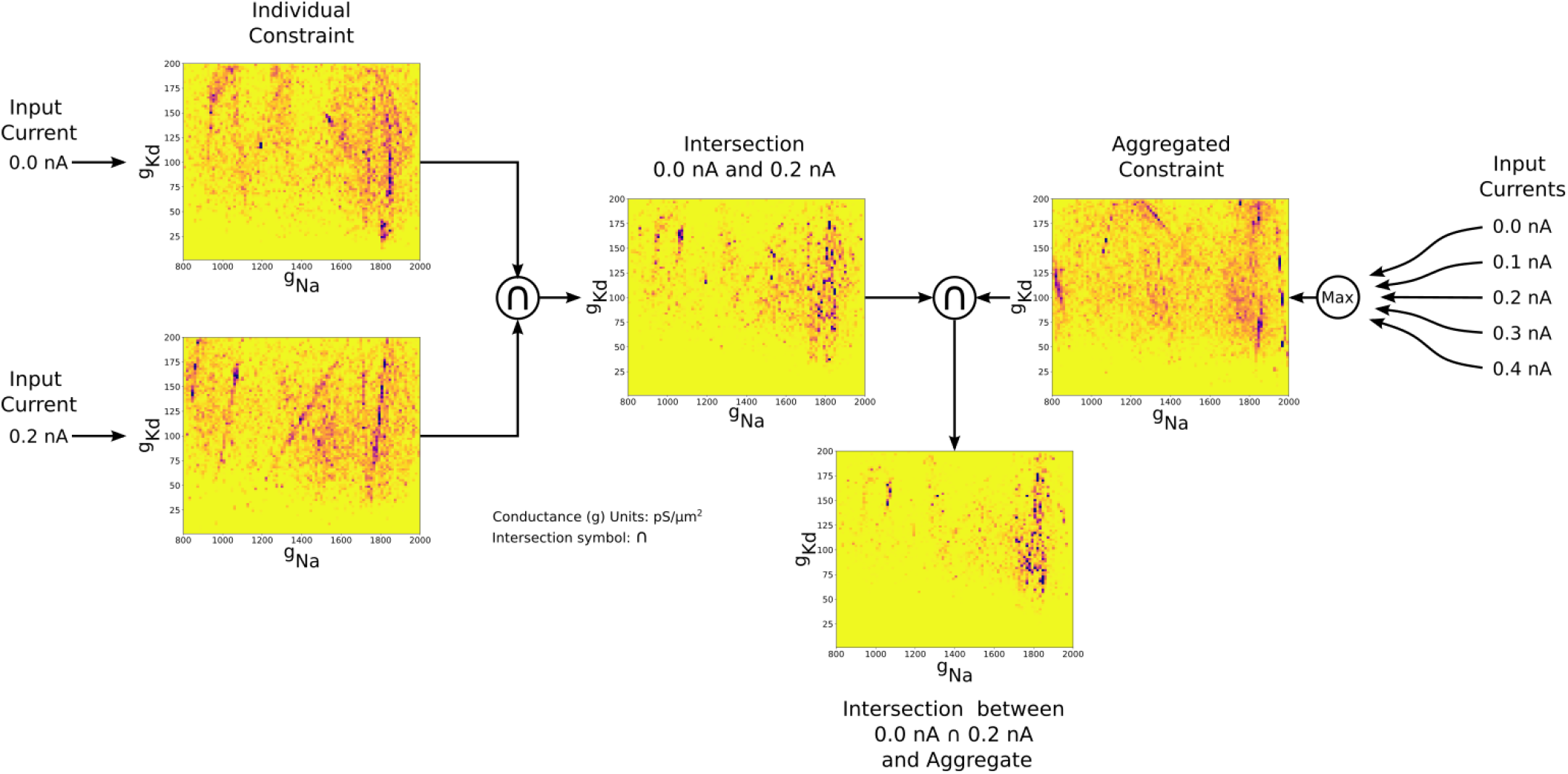
Two-dimensional (*g_Na_* and *g_Kd_*) kernel density estimates (KDEs) of the solution maps of different cases, for visualization of the relationship between the aggregate constraint and the individual constraints. The arrows indicate the flow of the operation. The KDEs of the 0.0 nA and 0.2 nA individual constraints are on the left. The KDE of the aggregate constraint is shown on the right. The intersection is the elementwise multiplication of two or more KDEs. The final intersection between 0.0 nA, 0.2 nA, and aggregate constraint shows the contribution of the 0.0 and 0.2 nA individual constraints to the aggregate constraint. The colormap scales have been adjusted so that values are visible.

One major result flows from this analysis. The distinctive features (lines, clumps,…) found in the individual constraints are found in the intersection even though they were indiscernible in the aggregate constraint results. This underlines the efficacy of the MCMC algorithm in solving for a solution map under the aggregate constraint.

In the next section we formalize these intersections using the Wasserstein distance metric.

#### AM model – Distances between posterior distributions

To demonstrate the differences between the posterior distributions of the AM model between an individual constraint and the aggregate constraint, we computed the approximate Wasserstein distance for each individual injected current’s posterior distribution taking into account all nine parameters as well as individual parameters against the aggregate constraint’s posterior distribution. The approximate Wasserstein distance provides a quantitative metric of the total cost required to transform one probability distribution to another probability distribution (***Givens and Shortt, 1984***). The results are displayed in ***Table 2***, with the rightmost column showing the Wasserstein distance of the entire posterior distribution (all nine parameters used) for a specific injected current. Overall, the posterior distribution of no injected current (0.0 nA) yields the closest posterior distribution to the aggregate constraint, followed by 0.2 nA, then 0.1 and 0.3 nA.

**Table 2.**
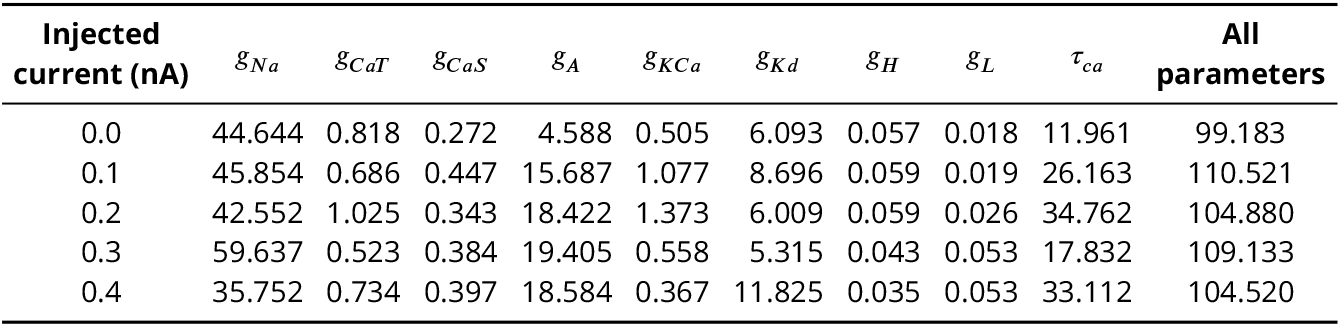
Wasserstein distances for each posterior (individual parameters and all parameters) of individually sampled injected currents against the aggregated constraint’s posterior.

The parameter’s posterior distributions that are the most different between an individual constraint and the aggregate constraint are the distributions for the parameter *g_Na_* and to a lesser extent for *g_A_*, *g_Kd_*, and *τ_Ca_*. This observation for the Wasserstein distances is consistent with the 1D marginal solution maps of the posterior when comparing the aggregate constraint in ***Figure 3*** with ***Figure 2*** and the additional solution maps of the posterior for other individual currents in the Appendix.

## Discussion

***Alonso and Marder*** (***2019***) presented a complex and realistic neurological model and proposed visualization techniques in order to help understand how different parameter sets can have similar model outputs. They identified the challenge to find multiple parameter sets that are optimal in the sense that they generate good fits between data and model outputs. Consequently, this raises the need to develop algorithmic approaches in order to find these multiple sets. In the present work we design the inverse problem in a Bayesian framework, where multiple optimal parameters are part of a single multimodal posterior distribution. Furthermore, we utilize parallel tempering MCMC in order to recover the multiple modes in the posterior and visualize them in the solution maps. As a result, we have filled an important need in the field of neuroscience.

### Interpreting solution maps and choosing parameter sets

The solution space that the MCMC algorithm provides can be overwhelming especially when one is familiar with the classical gradient approach to an inverse problem (i.e., a single solution). Here, the algorithm instead provides a space of solutions from which one can pick a solution.

#### Single-Solution Approach

As detailed in the Results, a parameter set exists that will produce the lowest loss value for each of the individual and aggregate constraints (see ***Table 1***). Many sets of solutions also will provide extremely low loss values. For instance, we found, for the solution space for the injected current of 0.0 nA, a total of 889 solutions within 0.5% of the best solution and a total of 1,595 solutions with 1% of the best solution (see ***Table 3***).

**Table 3.** Number of solutions given the injected current as a function of percentage difference from the best solution. For instance, for a 0.0 nA injected current, 889 solutions are 0.5% away from the best 0.0 nA solution. MCMC is able to find these multiple acceptable solutions. ***Figure 6*** visualizes an example set of voltage traces generated from these solution parameter sets.

| Injected current (nA) | 0.5% | 1.0% |
| --- | --- | --- |
| 0.0 | 889 | 1595 |
| 0.1 | 916 | 1618 |
| 0.2 | 21 | 118 |
| 0.3 | 120 | 233 |
| 0.4 | 149 | 265 |

Even though the loss values are small for many parameter sets, the voltage traces produced by these sets are different. To illustrate these differences, in ***Figure 6*** we plotted the first five voltage traces in order of the loss value from best to worse with respect to the ground truth shown at the top in orange. The change in loss value is smaller than 10^−2^ and nearly identical for the last two traces d and e. These traces are visibly different, however, thus illustrating that MCMC is able to produce different parameter sets for a nearly identical loss value (posterior distribution). This result highlights how this ODE neuron model can produce multiple solutions as shown by ***Alonso*** ***and Marder*** (***2019***).

#### Multimodal Approach

Many more nearly-optimally fitting solutions exist, however. One can look for multiple modes or sets of possible solution regions: this is illustrated in the 1D and 2D marginals described in the Results. In the 1D marginals one can quickly gauge the important values within a range by the different peaks observed within the distribution as well as its sensitivity by the width/spread under peaks. Furthermore, the size of the range can be determined if distributions appear to abruptly stop at the boundaries. For instance, in the 1D marginal for parameter *g_KCa_* found in ***Figure 2***, it appears that the range was too small as the distribution begins a descent at the upper bound. Such insight can lead the modeler to reconsider the size of the range of the parameters. The 2D Marginals provide a fast method to determine which parameters are highly correlated. Taken together, one can begin to look at solution areas that previously were left unexplored, perhaps opening new avenues for investigation.

### Impact of loss function on recovered parameters

The impact of the choice of loss function becomes apparent in ***Figure 5***. There we see the ground truth voltage trace generated from the AM model with the ground truth set of parameters (orange color). This trace serves as the data in our inverse problem. To measure the fit between data and model outputs, we consider a particular loss function that is derived from the loss function employed by ***Alonso and Marder*** (***2019***). After running the MCMC algorithm to recover the parameters of the model, we select the set of parameters with the lowest loss value and plug them into the model to generate the voltage traces, shown in blue color in ***Figure 5***. Comparing the orange (ground truth) and the blue trace graphs, we observe discrepancies between the traces that may appear as inadequate fits from a physiological perspective. These discrepancies are caused by the loss function, which defines a distance metric between traces. In essence, a possible limitation of the chosen loss function is that it quantifies disparate voltage traces as too similar by giving it a low loss value. This demonstrates the difficulty of finding appropriate losses for inverse modeling in neuroscience and is one main reason for manual and heuristic approaches for solving inverse problems governed by neural dynamics models.

**Figure 5.**
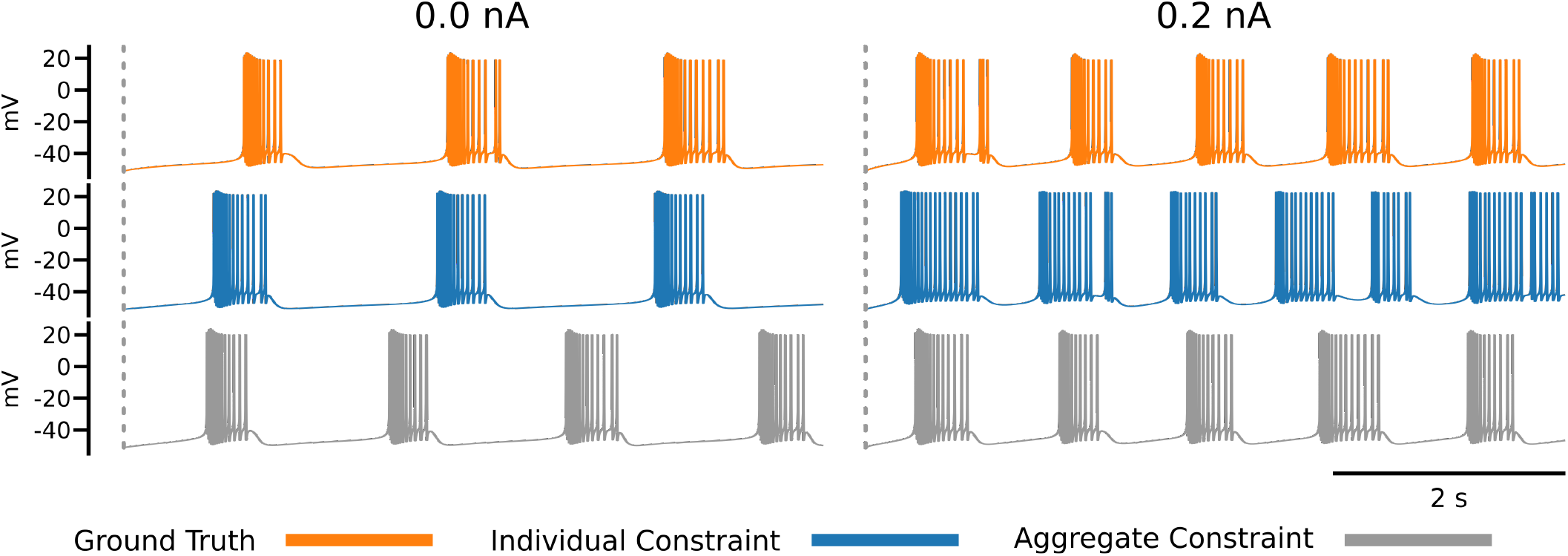
Each row of this figure shows the voltage traces for the ground truth (top orange), for the best solution sets given the individual constraints (middle blue) and the best solution set given the aggregate constraint (bottom grey). Each column shows the traces for different injected currents. Loss values for these parameter solution sets are 0.0008 for the 0.0 nA individual constraint, 0.0348 for the 0.2 nA individual constraint, and 8.8109 for the aggregate constraint.

**Figure 6.**
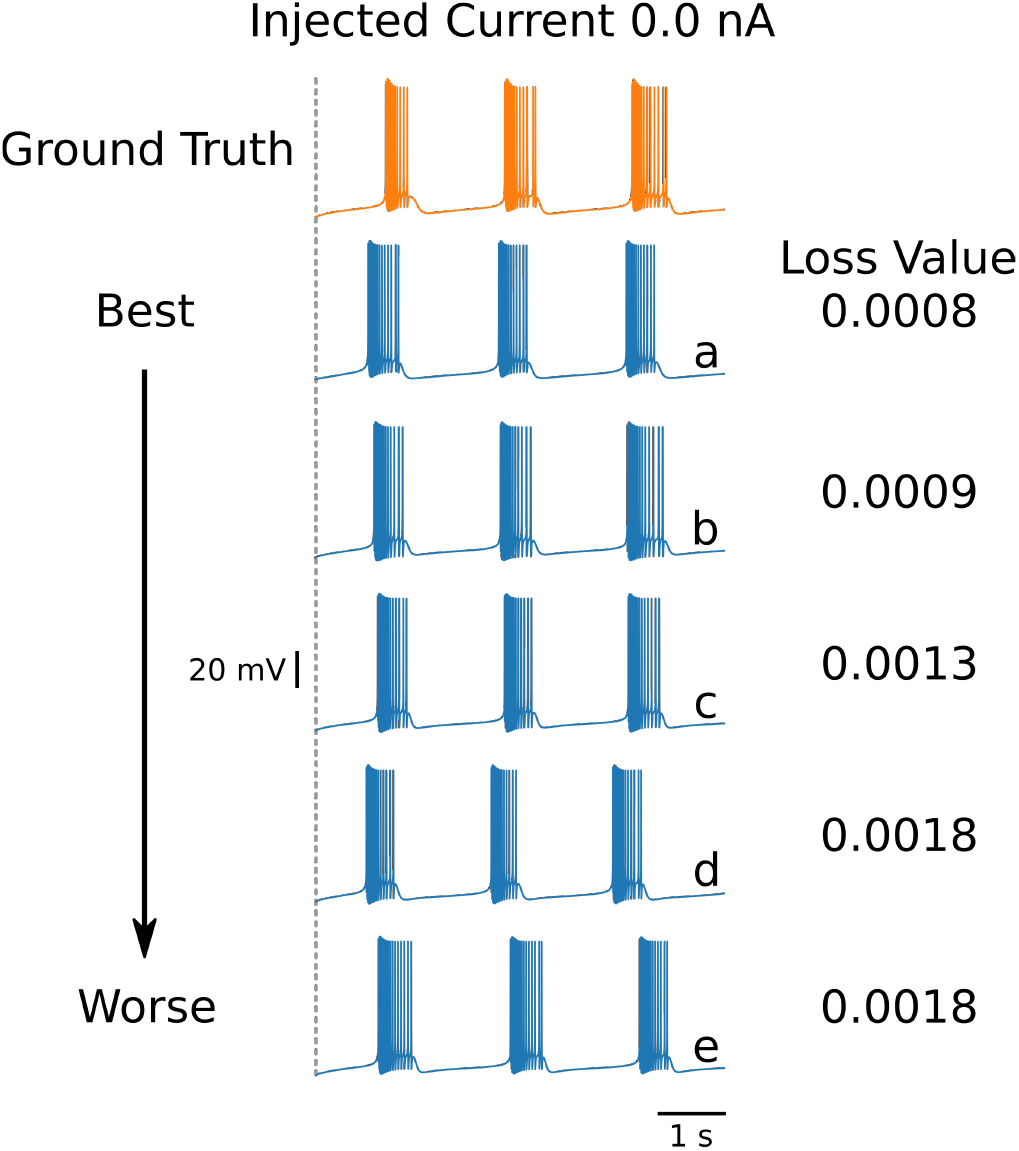
First 5 solutions for the injected current of 0 nA. These solutions are ranked from “best” to “worst” (a–e) according to their loss. Even a nearly identical loss can produce a qualitatively different voltage trace, as shown by traces d and e

To address this issue, ***Alonso and Marder*** (***2019***) invested efforts to refine the loss to specific inference setups that are targeted. This approach can be prone to human bias and is labor intensive. An alternative approach is taken in the present work because we aim at automated and algorithmically guided techniques. The idea is to augment the measure for fitting data and model outputs with additional information. Here we propose to augment the loss function with additional voltage traces that are generated at different input currents. This approach results in the inverse problem setup with aggregated current constraint, and we observe better-behaved solutions from an inference perspective, shown as the gray voltage traces in ***Figure 5***. An alternative approach to the aggregate current constraint would be to design new loss functions (for individual currents), which is out of the scope of this work.

### Physiological meaning

Extracting physiological information from the solution maps is the ultimate goal of this exercise. For instance, one can quickly rank the importance of a parameter by looking at the 2D marginals. Looking at the bottom row of ***Figure 2***, we observe that *τ_Ca_* is highly correlated to *g_Kd_* while *g_CaT_* and *g_CaS_* are more independent of *τ_Ca_* for the injection current constraint 0.0 nA. Therefore this MCMC analysis shows that the calcium component of the afterhyperpolarization (AHP) needed to replicate the ground truth spiking traces is dominated by the time constant *τ_Ca_*. To be able to reach such a conclusion with traditional optimization techniques would require a sensitivity analysis.

Alonso and Marder showed that multiple solutions exist for solving for the same neural outcome. MCMC complements this finding by showing that a much larger solution set can be found. We believe that this multimodal solution set is the norm rather than the exception. As models increase in complexity and therefore increase in parameters, the likelihood of a multimodal solution will increase. For instance, although studies of variability of neuronal behavior have concentrated on the role of ion channel density (e.g., ***Marder and Goaillard*** (***2006***)), changes in the voltage dependence of channels resulting from changes in phosphorylation (***Park et al., 2006***) or the binding of accessory proteins (***Bosch et al., 2015***) are likely to be equally important. The power of Bayesian inference together with MCMC is that these additional variables can be easily taken into account.

MCMC sampling operates within the bounds based on our prior assumptions such as our general knowledge, literature review, or colleagues. The MCMC algorithm “tests” these prior assumptions based on data and model by looking at the shape of the posterior, as mentioned above. Peaks at the boundaries of the ranges are likely to indicate too narrow a range of sampled parameter values (see parameters *g_A_* and *g_KCa_* in ***Figure 2***). This could lead experimentalists to look beyond their prior assumptions.

## Methods

### Details of the AM model

For the simulations governed by the AM model in Results, the bursting neuron ODE system from ***Alonso and Marder*** (***2019***) was used. The parameters used for the ground truth were from Model A of ***Alonso and Marder*** (***2019***) for consistency (conductances in uS, time constants in ms): *g_Na_* = 1076.392, *g_CaT_* = 6.4056, *g_CaS_* = 10.048, *g_A_* = 8.0384, *g_KCa_* = 17.584, *g_Kd_* = 124.0928, *g_H_* = 0.11304, *g_L_* = 0.17584, and *τ_Ca_* = 653.5.

The BDF and LSODA solvers for ODEs from the SciPy library (***SciPy 1.0 Contributors et al., 2020***; ***Petzold, 1983***) were utilized for the stiff ODE systems of the AM model. BDF and LSODA were used because of their overall convergence performance compared with that of other ODE solvers for stiff biological ODE models. LSODA has been shown to perform better in biological systems than many modern ODE solvers (***Städter et al., 2021***). The voltage trace generated by the ground truth parameters is shown in ***Figure 5*** in orange color.

### Loss function for measuring fit between data and model outputs

The main loss function used in the present study involving the AM model is the loss in Equation 4 in ***Alonso and Marder*** (***2019***), where we use the following coefficients: *α* = 1000, *β* = 1000, and *γ* = *δ* = *v;* = 0. This design of the loss has been highly informed by domain expertise, because it targets specific properties of voltage traces. We purposefully eliminate several components of the loss (i.e., *γ* = *δ* = *v* = 0). Rather than separating between bursters and non-bursters, our study uses the same loss function with fixed coefficient values throughout all experiments, because this allows us to compare inference results and it avoids possible human bias. Our approach is different from ***Alonso and Marder*** (***2019***), where the coefficients in the loss are adjusted depending on the inference setup. Our goal is to develop inference methods that can be carried out in an automated fashion. Here we focus on the two components of the loss function that we observed to be dominant, associated with *α*, the burst frequency mismatch, and with *β*, the duty cycle mismatch. Our objective is to test the multimodality of the inverse problem with parallel tempering MCMC methods.

### Loss function for individual and aggregate constraint

We use the loss function (detailed in Loss function for measuring fit between data and model outputs) to solve inverse problems, where one particular injected current is present; these studies are referred to as the individual current constraint. The individual current constraint (or “individual constraint” in short) is when the loss function value for a single current injection value (0.0 nA, 0.1 nA,…) is used for measuring the fit between data and model outputs.

In addition, we carry out experiments with multiple injected currents, referred to as the aggregate current constraint. The aggregate current constraint (or “aggregate constraint” in short) is when the maximal loss function value for the entire set of injected currents ({0.0, 0.1, 0.2, 0.3, 0.4} nA) is used for the misfit between data and model outputs. The aggregate current constraint is motivated by its similarity to fitting against a frequency-current (F-I) curve (***Hultborn and Pierrot-Deseilligny, 1979***).

### Introduction and background to MCMC

In this work we choose the Bayesian formulation (also called Bayesian framework), which is centered around Bayes’ theorem:

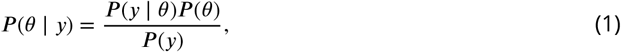

where *θ* denotes the parameters of the ODE model and *y* is the data. On the left-hand side of (1), *P* (*θ* | *y*) is the unknown *posterior* (i.e., conditional probability for the model parameters *θ* given the data *y*) On the right-hand side of (1), *P* (*y* | *θ*) is the known but often intractable *likelihood* (i.e., conditional probability for the data *y* given the model parameters *θ*). The likelihood term is responsible for measuring the fit between data and model outputs. Hence the loss function, which we detailed above, is evaluated each time the likelihood is computed. Further on the right-hand side, *P* (*θ*) is the known and typically tractable *prior* (i.e., probability of the model parameters *θ*). The term *P* (*y*) is called *evidence* (i.e., total probability of the data *y*); and since it is constant with respect to *θ*, it simply scales the posterior by a constant and can in practice remain unknown.

The first goal of MCMC algorithms, such as the Metropolis–Hastings algorithm (***Metropolis et al., 1953***; ***Hastings, 1970***), is to find the most likely model parameters, *θ*, that will produce the greatest posterior (the highest probability of the model parameters to reproduce the data). The second goal is to quantify parameter uncertainties. To this end, the algorithm visits smaller locations in the solution map of the posterior (i.e., lower probabilities of the model parameters to reproduce the data). However, these visits to lower-probability locations of the posterior happen adaptively and automatically at smaller frequencies. The concept of MCMC visiting locations of high probability at greater frequencies and locations of lower probability at smaller frequencies is a key property of the algorithm.

In detail, at each iteration of MCMC, the algorithm first proposes a new value for (one or multiple) parameters, *θ*_proposal_, by randomly choosing from a proposal distribution (e.g., a normal distribution centered at the value of the previous iteration *θ*_previous_). The likelihood *P* (*y* | *θ*_proposal_) and prior *P* (*θ*_proposal_) are evaluated at the proposed value and multiplied to obtain a new value of the posterior. In the second MCMC iteration, the algorithm includes the proposed *θ*_proposal_ in a sequence of visited points or discards it, where the aim is to more frequently keep points of higher probability. To determine whether *θ*_proposal_ is accepted or rejected relative to the previous *θ*_previous_, the algorithm takes the ratio of the proposed posterior to the current posterior. If the ratio is greater than one, *θ*_proposal_ must have a higher probability, and therefore this new parameter is saved because it is producing a better outcome. If the ratio is below one, then the algorithm applies a randomized rejection criterion such that some *θ*_proposal_ is stored at lower frequencies. Subsequently, this cycle repeats, where *θ*_proposal_ becomes the new *θ*_previous_, if acceptance was successful. Note that in this algorithm, dividing the new value by the previous value of the posterior, *P* (*y*) cancels out, thus showing that constant scaling factors are irrelevant for MCMC.

The Metropolis–Hastings algorithm constructs a sequence, also called a chain, of parameters (also called samples) that it visits during the iterations. This method is suitable for unimodal posterior distributions. However, it is not designed to capture multimodal posteriors (i.e., solution maps of the posterior with multiple peaks). Therefore, extensions of MCMC have been developed to deal with multimodal distributions. Early work by ***Marinari and Parisi*** (***1992***); ***Geyer and Thompson***(***1995***) employed simulated tempering, which is akin to simulated annealing algorithms from the field of optimization. More recently, parallel tempering MCMC (***Łącki and Miasojedow, 2016***; ***Vousden et al., 2016***) was proposed. This builds on ideas from simulated tempering and, additionally, uses multiple chains of single-chain methods, such as Metropolis–Hastings, in parallel. The key idea of tempering, which enables MCMC to discover multimodalities in posteriors, is that the posterior is taken to a power with an exponent, *γ*, between zero and one: 0 < *γ* < 1. Hence we obtain *P ^γ^* (*θ* | *y*). This has the effect that smaller locations in the solution map of the posterior are elevated and can be visited more frequently by Metropolis–Hastings.

### MCMC method for estimating parameters in AM model

This section provided details about our implementation for estimating parameters and their uncertainties in AM models. Our AM model and MCMC methods constitute the following components:

- Neuron model component: Computes voltage traces for the AM model and a given set of candidate parameters
- Likelihood component: Computes the Bayesian likelihood using the loss function based on the voltage trace obtained by neuron model component
- Prior component: Evaluates the prior based on the candidate set of parameters
- Estimator component: Proposes candidate parameters at random (the proposal sampler)

The sampler used was the adaptive parallel tempering sampler (***Miasojedow et al., 2013***) within the PyPESTO library (***Stapor et al., 2018***). Parallel tempering runs independent Markov chains at various temperatures and performs swaps between the chains. Each individual chain runs adaptive Metropolis–Hastings MCMC.

The loss functions for individual and aggregate constraints are evaluated for each candidate parameter in the MCMC algorithm within the likelihood term. The prior term is defined as a uniform distribution (i.e., unbiased prior) within predetermined boundaries, which were chosen sufficiently wide to permit physiologically relevant model parameters.

### Experimental setup

To investigate the robustness of parallel tempering MCMC on the inverse problem governed by the AM model, we use model A from ***Alonso and Marder*** (***2019***), which is defined by the parameters that are listed in Details of the AM model, as ground truth data. We use this model to generate five voltage traces each using different injected currents: 0.0, 0.1, 0.2, 0.3, and 0.4 nA. Noise to simulate physical measurement noise was not added to the voltage trace of the ground truth, for several reasons. To have an experimental setup similar to ***Alonso and Marder*** (***2019***), we also did not alter the ground truth trace data. Furthermore, MCMC methods are typically not detrimentally affected by noisy data. Further, the Bayesian framework together with MCMC as a solution algorithm allows noise to be modeled using an additional parameter (and prior) within the inverse problem. This is a possible direction for future research.

The conductances for the ion channels and the time constant for the calcium channel are the nine parameters that we target for inference. Each inference with individual current constraint consisting of running the MCMC algorithm for a predetermined number of iterations/samples. Additionally, for the inference with aggregate current constraint, a single run of MCMC was performed. Each MCMC run used 24 to 32 parallel chains and 10,000 to 14,000 samples per chain. Geweke’s convergence diagnostic was used to determine the burn-in of each chain (***Stapor et al., 2018***). The burn-in is an initial phase of MCMC where the gathered samples do not satisfy certain statistical properties to be considered adequate samples of the posterior. In certain cases, despite parallel tempering, there are MCMC chains that can remain static through the parameter space. Results from chains that did not move sufficiently (ratio of standard deviation to mean) were filtered out in postprocessing.

### Software improvements for accelerating parallel tempering MCMC

We reduced the computational time of the MCMC inference by carrying out various optimizations of the source code. Increasing the number of Markov chains benefits the detection and exploration of multimodal posteriors; however, it also increases the computation time. To shorten the computation time, we parallelized the MCMC algorithm to run on each CPU core concurrently using multiprocessing packages (***conda-forge community, 2015***). This approach significantly decreased computation time because of the parallel nature of the chains. The total runtime for each MCMC run was between 24 and 48 hours. We utilized three different machine architectures: (1) the Broad-well nodes (36-core, 128 GB RAM) on the parallel cluster Bebop at Argonne National Laboratory, (2) a workstation equipped with a dual 16-core Intel Xeon 4216 CPU (total of 32 cores and 64 threads) with 128 GB of RAM, and (3) a workstation with a 32-core (64-thread) AMD Ryzen 3970x CPU with 128 GB of RAM.

## Acknowledgments

We gratefully acknowledge the computing resources provided on Bebop, a high-performance computing cluster operated by the Laboratory Computing Resource Center at Argonne National Laboratory.

The efforts of M.C., C.J.H., R.K.P, and N.S. was supported in part by the NIH R01 NS109552. The effort of G.I. was supported by the NSF Mathematical Sciences Graduate Internship. The effort of J.R. is based in part upon work supported by the U.S. Department of Energy, Office of Science, under contract DE-AC02-06CH11357. The effort of Y.C.W. was supported in part by the Center for Energy and Sustainability at California State University, Los Angeles, and the National Science Foundation under Award No. HRD-1547723.

## Appendix 1 MCMC solution maps of posteriors for individual constraints

**Appendix 1 Figure 1.**
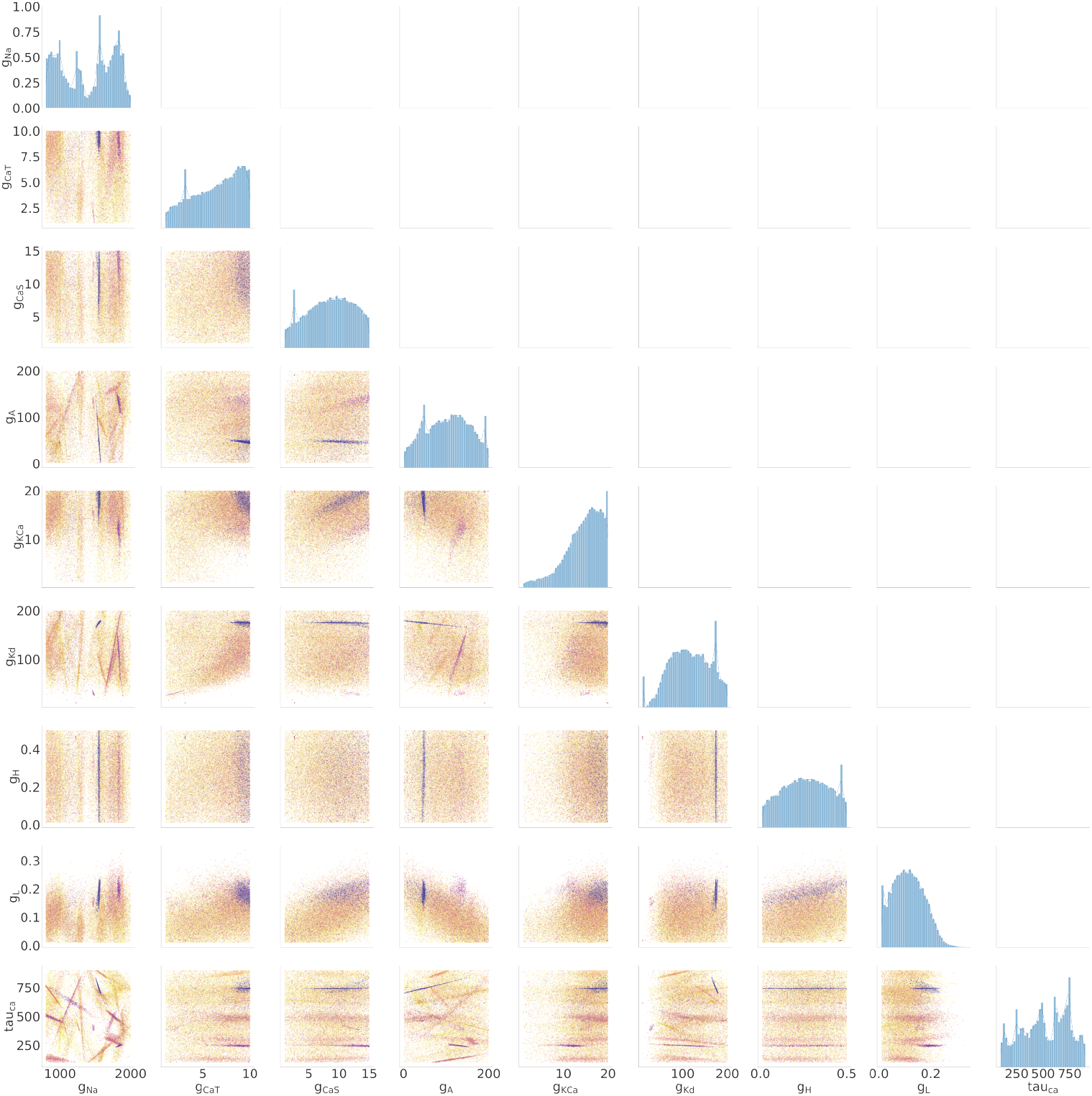
*0.1 nA results*: Diagonal shows the 1D marginals for each of the parameters. The remaining plots show the 2D marginal relating one parameter with an other. The color represents the loss value from blue to yellow (low to high). The density is represented by the opacity (i.e., light to dark shows low to high densities). Each point is set to a transparency level of *α* = 1*e* − 3 where *α* = 1 is opaque. As such, darker regions have more dots and indicated higher probabilities for good fits between data and model outputs.

**Appendix 1 Figure 2.**
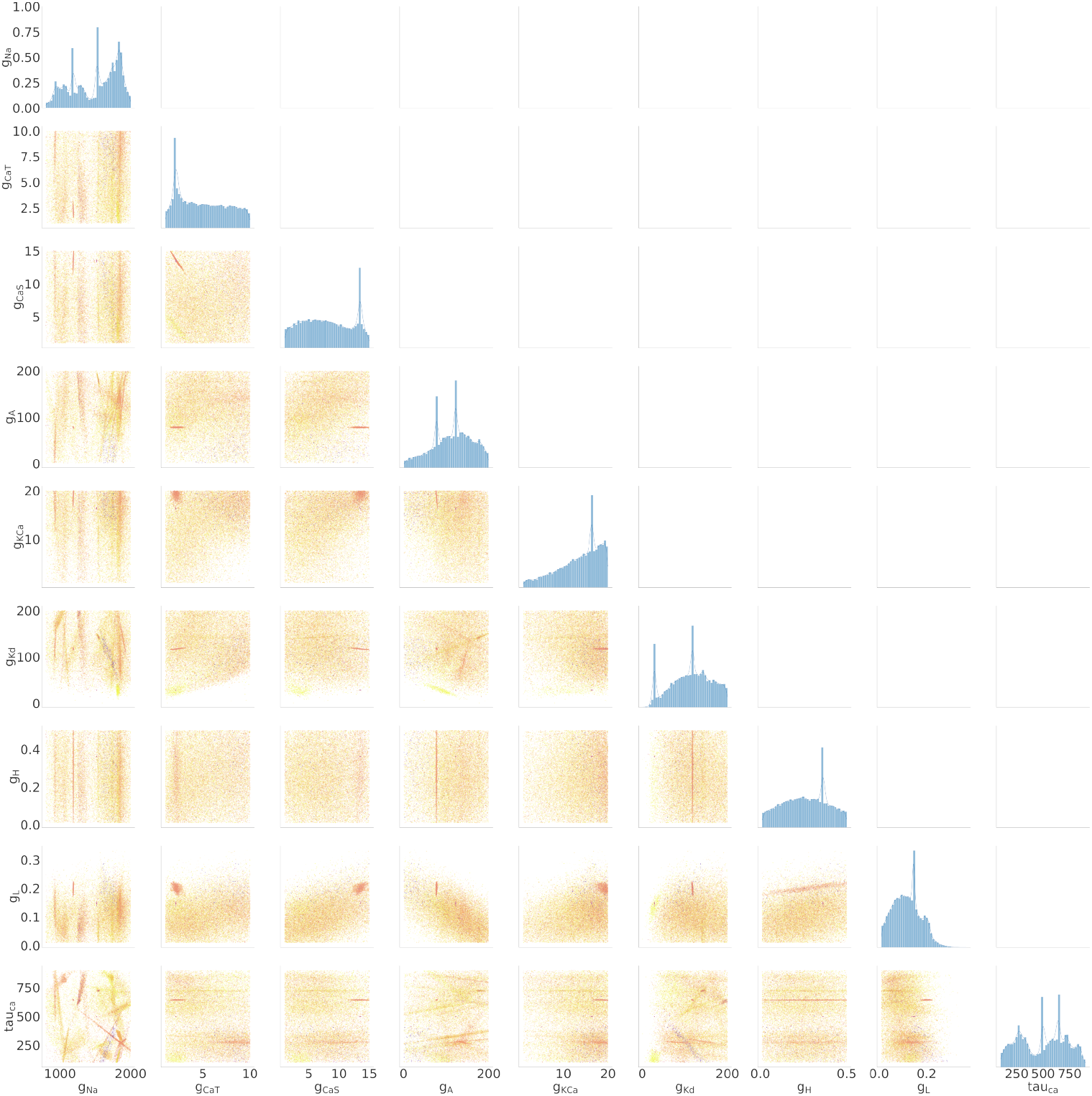
*0.2 nA results*: Diagonal shows the 1D marginals for each of the parameters. The remaining plots show the 2D marginal relating one parameter with an other. The color represents the loss value from blue to yellow (low to high). The density is represented by the opacity (i.e., light to dark shows low to high densities). Each point is set to a transparency level of *α* = 1*e* − 3 where *α* = 1 is opaque. As such, darker regions have more dots and indicated higher probabilities for good fits between data and model outputs.

**Appendix 1 Figure 3.**
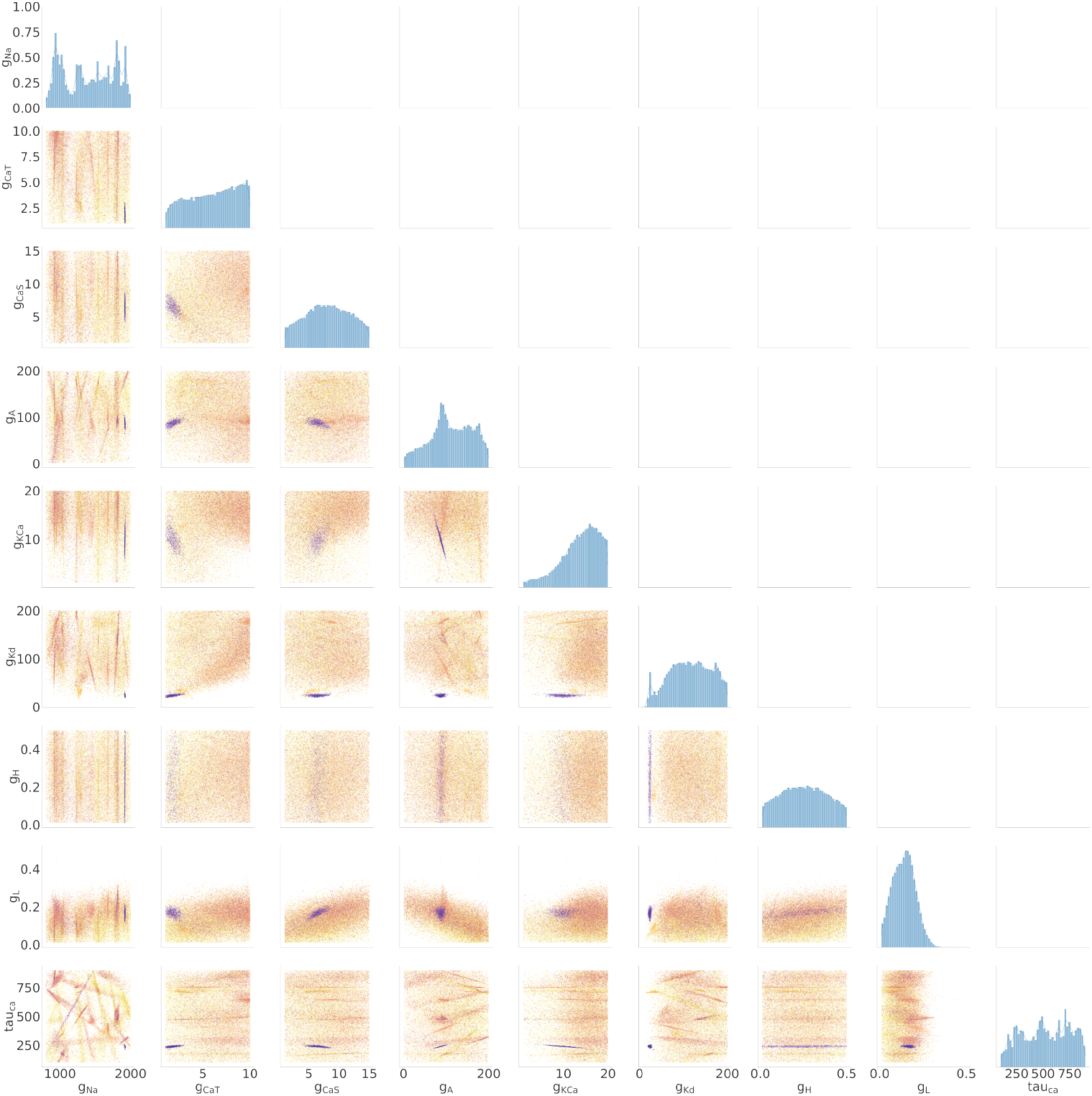
*0.3 nA results*: Diagonal shows the 1D marginals for each of the parameters. The remaining plots show the 2D marginal relating one parameter with an other. The color represents the loss value from blue to yellow (low to high). The density is represented by the opacity (i.e., light to dark shows low to high densities). Each point is set to a transparency level of *α* = 1*e* − 3 where *α* = 1 is opaque. As such, darker regions have more dots and indicated higher probabilities for good fits between data and model outputs.

**Appendix 1 Figure 4.**
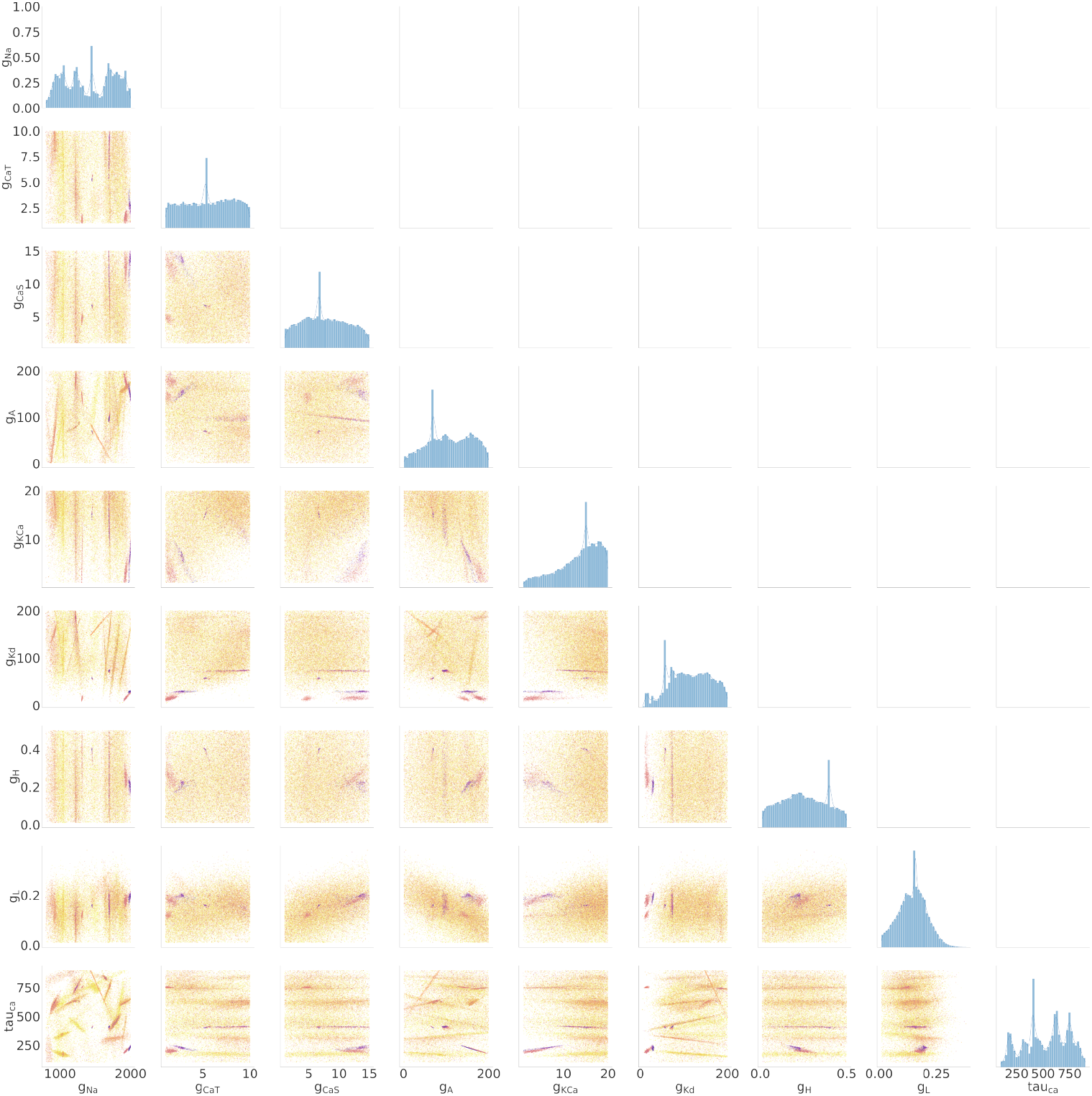
*0.4 nA results*: Diagonal shows the 1D marginals for each of the parameters. The remaining plots show the 2D marginal relating one parameter with an other. The color represents the loss value from blue to yellow (low to high). The density is represented by the opacity (i.e., light to dark shows low to high densities). Each point is set to a transparency level of *α* = 1*e* − 3 where *α* = 1 is opaque. As such, darker regions have more dots and indicated higher probabilities for good fits between data and model outputs.

